# The ability of bumblebees *Bombus terrestris* (Hymenoptera: Apidae) to detect floral humidity is dependent upon environmental humidity

**DOI:** 10.1101/2021.08.13.456254

**Authors:** Amy S. Harrison, Sean A. Rands

## Abstract

Flowers produce local humidity that is often greater than that of the surrounding environment, and studies have shown that insect pollinators may be able to use this humidity difference to locate and identify suitable flowers. However, environmental humidity is highly heterogeneous, and is likely to affect the detectability of floral humidity, potentially constraining the contexts in which it can be used as a salient communication pathway between plants and their pollinators. In this study, we use differential conditioning techniques on bumblebees *Bombus terrestris audax* (Harris) to explore the detectability of an elevated floral humidity signal when presented against different levels of environmental noise. Artificial flowers were constructed that could be either dry or humid, and individual bumblebees were presented with consistent rewards in either the humid or dry flowers presented in an environment with four levels of constant humidity, ranging from low (∼20% RH) to highly saturated (∼95% RH). Ability to learn was dependent upon both the rewarding flower type and the environment: the bumblebees were able to learn rewarding dry flowers in all environments, but their ability to learn humid rewarding flowers was dependent on the environmental humidity, and they were unable to learn humid rewarding flowers when the environment was highly saturated. This suggests that floral humidity might be masked from bumblebees in humid environments, suggesting that it may be a more useful signal to insect pollinators in arid environments.

## INTRODUCTION

Most plant-pollinator interactions involve the plant producing signals that manipulate the pollinator in order to bring it to the plant. These signals can work across many different sensory modalities, and may include colour (Gumbert et al. 1999, Glover and Whitney 2010, Dyer et al. 2012), shape and pattern (Leonard and Papaj 2011, Lawson et al. 2017a), symmetry (Rodríguez et al. 2004, Krishna and Keasar 2018), scent (Miyake et al. 1998, Theis 2006, Raguso 2008, Lawson et al. 2018), temperature (Dyer et al. 2006, Whitney et al. 2008, Harrap et al. 2017), texture (Kevan and Lane 1985, Goyret and Raguso 2006, Whitney et al. 2009, Goyret 2010), electrostatic charge (Clarke et al. 2013, 2017). Recently, humidity has been suggested to be another sensory modality which could be used for signalling to pollinators. There is good evidence that flowers produce humidity that is elevated when compared to the immediate environment, having been demonstrated to occur in a number of flower species (Corbet et al. 1979, von Arx et al. 2012, Nordström et al. 2017, Harrap et al. 2020). Floral humidity is created by a combination of nectar evaporation and floral transpiration (Corbet et al. 1979, Azad et al. 2007, von Arx et al. 2012, Harrap et al. 2020) although the contribution of these two influences may vary between species, and appears to be different in its intensity from the humidity produced by non-floral vegetation (Harrap and Rands 2021). A sample of the floral headspace of 42 species found that 30 (71%) produce floral humidity of an intensity greater than would be expected from any conflating environmental humidity sources (Harrap et al. 2020) (such as the minimal humidity differences due to uneven air mixing in the sampling room, or humidity produced by water within the capped horticultural tubes that flowers were mounted in during sampling). The intensity of floral humidity produced by flowers, represented the average peak difference in relative humidity in the flower species’ headspace, compared to the background, reached up to 3.71% (in *Calystegia sylvatica*).

Floral humidity occurs widely and varies between species (Harrap et al. 2020), and does not appear to be limited to species visited by a particular group of pollinators (Harrap et al. 2020). Elevated floral humidity intensity has been observed in flowers pollinated primarily by moths (von Arx et al. 2012), flies (Nordström et al. 2017) and bees (Corbet et al. 1979). Sensitivity to environmental (non-floral) humidity is well reported in insects (Havukkala and Kennedy 1984, McCall and Primack 1992, Kwon and Saeed 2003, Peat and Goulson 2005, Liu et al. 2007, Lin et al. 2014, Enjin 2017), and it is possible that these variations in floral humidity could function as a foraging cue for pollinators, especially since many different taxa of insect have been shown to be able to detect humidity differences similar to those seen between flowers and their surrounding environment (Harrap et al. 2021). Evidence for floral humidity acting as a signal currently comes from two pollinator species. The hawkmoth *Hyles lineata* was demonstrated (von Arx et al. 2012, von Arx 2013) to show a preference to artificial flowers producing floral humidity comparable to that produced by *Oenothera caespitosa* (which is naturally pollinated by *H. lineata*), and the bumblebee *Bombus terrestris* is capable of learning and differentiating between artificial flowers that mimic natural patterns (Harrap et al. 2021). Other existing evidence of the capacity of pollinators to respond to floral humidity is limited (von Arx 2013), with non-experimental observations that flies may use floral humidity in addition to other floral display traits produced within Indian alpine environments (Nordström et al. 2017). Overall, given that floral humidity is present in many flower species, and can potentially be detected by insect visitors, it is therefore likely that floral humidity will provide at least one component of the complex multisensory advertisement that a flower produces to attract its pollinating visitors (Raguso 2004, Leonard et al. 2011).

Signals that are more easily detectable due to contrast with the background will enhance the learning response and promote constancy (Spaethe et al. 2001, Dyer and Chittka 2004a, Heuschen et al. 2005, Chittka and Spaethe 2007). Pollination interactions exist against a background of noise due to the local climate (Wilson et al. 2015, Lawson et al. 2017b, Lawson and Rands 2019), which can reduce signal detectability and learning responses (Chittka et al. 1999, Dyer and Chittka 2004a, Kaczorowski et al. 2012, Lawson et al. 2017b). Environmental humidity is highly heterogenous (Lionello et al. 2006, Alarcón et al. 2008, Tichy and Kallina 2014, Enjin 2017) and is therefore likely to be a significant source of noise for floral humidity signals, but this effect has not yet been explored. If this is the case, environmental humidity may significantly affect the detectability of floral humidity, potentially constraining the contexts in which it can be used as a salient communication pathway between plants and their pollinators. In this study, we consider whether the ability of a pollinator to use floral humidity could be dependent upon the environment that the signal is presented within, which in turn tells us whether floral humidity is important regardless of environmental or weather conditions, or whether changes in the environment can impact on the effectiveness of communication using this sensory pathway.

In this study, we use differential conditioning techniques to explore the detectability of an elevated floral humidity signal comparable to natural flowers when presented to bumblebees *Bombus terrestris audax* (Harris) against different levels of signalling noise comparable with native conditions. This species is suitable for these experiments, as *B. terrestris* foragers have been used in similar differential conditioning experiments to those described here, showing a capacity to detect floral humidity and associate nectar with both humid and dry artificial flowers (Harrap et al. 2021). If there is no effect of noise on detectability, making floral humidity a robust signal, we would expect that the bees’ learning responses would not be affected by the environmental conditions.

## METHODS

This study used differential conditioning techniques to test the robustness of a constant floral humidity signal generated at 2% against ambient lab conditions against varying levels of background humidity. Signal detectability was determined by observing the learning response of foraging bees to a sucrose reward associated with relative humidity differences between humid and dry artificial flowers. Observations were made when each of the flower variants was associated with the reward, and the other was non-rewarding. Trials were repeated in Low, Ambient, Medium and High relative atmospheric humidity conditions to alter the contrast of humid flowers with both the background and with dry flowers.

### Ethical Information

This study did not require ethical approval or licensing. All experimental procedures and animal husbandry were conducted according to the ASAB/ABS (2006) guidelines for animal behaviour experiments.

### Bee colonies and flight arenas

Nestboxes of *B. terrestris* (either from Biobest, Westerlo, Belgium or Syngenta-Bioline, Clacton-on-Sea, UK) were housed in an internal laboratory maintained at 21°C and 40 ± 5% RH (relative humidity), under a 12 hour day-night cycle light ladder (Sylvania Activa 172 Professional 36 W fluorescent tubes, Havells-Sylvania Germany GmbH, Erlangen, Germany). The nestboxes were connected to separate 72 × 104 × 30cm flight arenas by gated tunnels where access could be controlled by the experimenter. The floor of each arena was covered with a layer of Advance Green Gaffer tape (Stage Electrics, Bristol, UK), and the lid of each was made of UV-transparent Perspex which allowed observations to be made externally. Each arena also had six sliding doors which could be opened to gain access to flight space. Outside of the trials, the nestbox had free access to the arena where pre-training stimulus flowers were present. These flowers lacked any signals that bees were tested on during learning trials, and were made from 60ml specimen jars (Sterilin PS, Thermo Fisher Scientific, Newport, UK) with white lids holding an upturned Eppendorf tub lid (Hamburg, Germany). Feeding flowers had slightly varying appearances to prevent bees developing preferences for a particular flower colour or feeding-well size and encourage exploration of new flowers. Once a day, flowers were filled with a 30% sucrose solution, and a further food source was provided in the form of a PCR rack filled to one third capacity with the sucrose solution. Two teaspoons of pollen were administered directly into the nests three times a week. Whilst bees were feeding in the arena outside of learning trials, individual foragers were marked using non-toxic paint of various colours to allow for visual discrimination of each bee.

### Artificial flowers: design and construction

The artificial flowers used were constructed according to the design of the ‘passive artificial flower’ used by Harrap et al. (2021). The flowers were identical except for their internal components, which were controlled to create a humid variant with elevated floral headspace humidity, and a dry variant. The main body of each flower consisted of a 60ml specimen jar and white lid (Sterilin, Newport). The lids were altered by cutting a 35mm hole in each, leaving the edges and inside threading intact so they could still be secured onto the jars. The transparent plastic body of the jars were coated in black electrical tape, which obscured the contents of the jars, preventing any visual learning by bees. To make the flowers distinguishable to the experimenter, half the jars were labelled with two two-digit odd numbers, and half with two two-digit even numbers in small white font positioned 180º from each other around the base of the jar. It was important that the landing surface of the flowers allowed for the different levels of humidity created by the internal components to be detected by visiting bees. Square segments of translucent gauze material (cut from Teresia curtains, Ikea, Leiden) were stretched over the jar and held modified lids so that the surface was taut. After screwing the lids in place, excess gauze was cut away to prevent any visual dissimilarities between flowers. Feeding wells for the flowers were made from detached 0.5ml Eppendorf tube lids, which were painted black and positioned centrally on the gauze surface during experiments.

The internal components of each flower consisted of three 40mm coloured sponge discs (cut from cellulose sponge wipes, Co-op, Manchester), with the top-most sponge always being green to achieve visual uniformity between all flowers. These sponges were stored in their original packaging until they were required to avoid additional drying out.

### Artificial flowers setup

For each trial, four flowers of each of the dry and humid variants (described below) were set up. Specimen jars were selected so that four evenly and four oddly numbered jars were used, each type associated with a flower variant. Whether odd jars were used for humid or dry flower variants alternated with each experiment.

For dry flower variants, sponge discs were cut out and placed directly into the flowers without further adjustments, followed by the securing of the gauze and lid. All lids were wiped with ethanol before being attached to remove any olfactory or chemical cues.

For humid variants, sponges were thoroughly soaked prior to the experiments in water that had been left to adjust to the room temperature overnight in a glass jug. Each disc was wrung out under the water before placing inside the flower to achieve full saturation, before a gauze sheet was stretched over the top and secured using a lid. If the gauze sheet got wet during preparation, the sheet was discarded and a dry sheet was used so that foraging preference would not be impacted by the presence of water on the surface.

In order to check that the humid variants created at least a 2% rise in relative humidity compared to background levels, a hand-operated hygrometer was used to read the relative humidity above each flower. Harrap et al. (2021) report that a more accurate reading of the relative humidity difference when the hand held hygrometer read 2% could be taken using DHT-22 humidity probes (Aosong Electronics Co., Ltd., Huangpu) attached to a 6-axis articulated *Staublie* RX 160 robot arm (Pfäffikon, Zurich), and was roughly 1.36%. This value reflects the differences in relative humidity observed by the same authors between various flower species and between flowers and background humidity, and is therefore a similar value to the change in relative humidity that might be experienced by a foraging bee in the natural environment. Flowers were de-assembled and sponges re-soaked if the hygrometer did not report a 2% RH rise. In the case of the dry flower variants, the RH value of the floral headspace was recorded at the start of an experiment using the hand-held hygrometer so that the humidity produced by the dry flowers could be monitored throughout the experiment.

It was also important to ensure that the two flower variants were not distinguishable by their temperature which may have occurred because of evaporation or the water temperature. A thermal imaging camera (FLIR Systems, Wilsonville) was used check the temperature of the landing surface of each flower. If either flower variant displayed a temperature difference of more than 1ºC, the warmer variant was placed into a 5ºC refrigerator for a few seconds. The flowers were then removed and accepted for use if the temperature difference was reduced to below 1ºC. Each flower then received an Eppendorf lid feeding well, which was loaded with either a 25ml drop of water or 30% sucrose solution as appropriate (see details of bee trials below).

### Artificial flower maintenance

During an experiment, artificial flowers were removed following the end of each foraging bout, when they were cleaned to remove any olfactory cues (Stout and Goulson 2001, Pearce et al. 2017) that might affect foraging decisions in subsequent bouts. Flowers were cleaned by removing the lids and discarding the gauze sheets. Each lid was wiped with ethanol using a cotton bud and dried using blue paper roll to prevent any additional humidity differences arising because of the ethanol. Each flower was fitted with an unused sheet of gauze, which was secured by replacing the lid and trimmed to remove any excess material. Feeding wells were cleaned using blue paper towel to prevent the build-up of any temperature differences that may have developed during the previous trial, and 25ml of fresh 30% sucrose solution or water was applied as appropriate. In the dry flower variants, the RH of the floral headspace was then measured using the hand-held hygrometer to check that the value had not fluctuated by 2% or more from the RH value observed during set-up the start of the experiment. Fluctuations may have arisen due to sponges gaining water from the surrounding environment during trials, especially in the Medium or High humidity environments. If the RH value of a flower was found to have fluctuated by 2% or more, then the lid and gauze were removed, and the sponges replaced with new dry sponges. Used spongers were discarded and the gauze and lid refitted. Furthermore, at the end of the foraging bout that contained the 35^th^ visit, all sponges in the dry variants were discarded and replaced with new dry sponges to minimise the risk of any water accumulating in the dry sponges. In the case of humid flowers, after cleaning and reassembly, RH was measured using the hand-held hygrometer. Flowers were de-assembled and sponges were re-soaked if the hygrometer did not report a rise of at least 2% against ambient humidity in the bee lab. The initial peak in humidity generated after assembly of humidity flowers has been shown to last for ten hours, meaning that for the duration of a learning experiment, the relative humidity difference between variants could be considered stable at roughly 2% (Harrap et al. 2021).

### Environment treatment setup

Bees were assigned to one of four environmental treatment groups designed to reflect the extremes and intermediate humidity conditions that may be experienced in the natural environment. The environment treatment groups were: *i*) Low humidity maintained at 20-25% RH, *ii*) Ambient humidity environment, maintained at 35-45% RH, dependent on the conditions in the laboratory, *iii)* Medium humidity environment where humidity was maintained at 67.5-72.5% RH, and *iv*) High humidity environment where humidity was maintained at 92.5-97.5% RH. The intensity of floral humidity produced by humid artificial flowers was the same in each environment, meaning that contrast with the background humidity would be 2% RH in the Ambient environment, but would be increased in the Low, decreased in the Medium, and decreased further still in the High. The method for setting up each treatment is described below.

#### Low humidity environment

Humidity in the arenas was lowered using a Manli Cordless Hair Dryer (Oblong Technology Company, Kowloon). Before the start of an experiment, the hairdryer was turned on inside the empty arena with two doors open to allow air flow. The handheld hygrometer was used to observe when the arena reached 20% RH. The temperature of the arena was recorded before drying, and again after drying. As the dryer recirculated unheated air only, the temperature inside the arena remained within 3ºC of the ambient arena temperature before drying. When 20% RH was achieved, the hair dryer was turned off and removed. Artificial flowers were introduced into the arena according to the test group and a learning trial was begun.

#### Ambient humidity environment

The RH of the arena in the Ambient environment group was not changed from background levels observed on the day of an experiment as relative humidity conditions inside the bee lab were controlled at roughly 40% RH, which fluctuated slightly due to changes in outside weather conditions. Therefore the parameters for the atmospheric humidity to be used in the Ambient environment group was set at within 5% of 40% RH. At the start of a trial, RH inside the arena was checked to ensure conditions were within these parameters.

#### Medium humidity environment

Humidity of the arena was elevated using a 0.5L Vicks Personal Humidifier (Kaz Consumer Products, Sheffield, UK) which was filled with water that had been left overnight to adjust to room temperature and placed inside the empty arena before the start of an experiment. The temperature of the arena was measured using the hand-held hygrometer, and the humidifier was turned on. The humidifier was turned off and removed from the arena once the arena had reached 72.5% RH (roughly five minutes). Temperature was monitored at this point to check that the arena temperature was within 2ºC of the ambient arena temperature, which was consistently reported given the use of room-temperature adjusted water in the humidifier. Water vapour created by the humidifier was allowed to completely settle before introducing the artificial flowers (roughly 5 minutes). This was done to allow full visibility to return for behavioural observations as water vapour plumes clouded the arena, and so that water vapour would not condense on the surfaces of artificial flowers, to avoid any further alterations in floral humidity being introduced or any water droplets on the surface of flowers that may affect foraging preference. The artificial flowers to be used that day were then placed inside the arena and the first foraging bout could begin.

#### High humidity environment

In the Humid environment treatment group, the relative humidity of the arena was raised using the Vicks Personal Humidifier as in the Medium humidity treatment. The humidifier was turned off and removed once the arena had reached 97.5% (roughly 10 minutes). Temperature was monitored at this point to check that the arena temperature was within 2ºC of the ambient arena temperature, which was consistently reported given the use of room-temperature adjusted water in the humidifier. As in the Medium humidity treatment, water vapour was allowed to completely settle. Artificial flowers were introduced according to the test group, and the first foraging bout was begun.

### Maintenance of environment treatment groups

Before each foraging bout (the full period where a bee was in the arena, from the moment it entered until the moment it left), the humidity treatments were checked and then maintained as follows.

#### Low humidity environment

Humidity was ‘topped up’ to 20% before the start of each foraging bout. Once the bee had returned to the nest, flowers were removed and for cleaning and refilling as described previously. Whilst the flowers were cleaned, the hair dryer was placed into the arena and turned on with two doors open to allow air flow. The hand-held hygrometer was used to monitor relative humidity and temperature. When humidity reached 20% RH, the dryer was removed, and temperature was checked to be within 2ºC of ambient temperature. The cleaned flowers were then returned to the arena to start the next foraging bout.

#### Ambient environment

Between each foraging bout, humidity inside the arena was checked using the hygrometer to check that the arena had not fluctuated outside of the 35-45% RH parameter before beginning the next foraging bout. As conditions inside the bee lab were stable, this was consistently reported.

#### Medium humidity environment

Humidity in the arena was ‘topped up’ to 72.5% RH before each foraging bout. After the flowers had been removed, the humidifier was returned to the arena and turned on whilst the flowers were being cleaned and refilled. The relative humidity and temperature were checked using the hand-held hygrometer. When humidity reached 72.5% RH, the humidifier was removed, and any water vapour was allowed to settle before reintroducing the flowers for the next foraging bout.

#### High humidity environment

At the end of a foraging bout humidity was ‘topped up’ to 97.5% RH. Whilst flowers were cleaned and refilled, the humidifier was placed inside the arena and turned on. The hand-held hygrometer was used to observe when humidity was at 97.5% RH, and temperature was checked to be within 2ºC of ambient temperature. The humidifier was then removed, and the water vapour was allowed to settle. The flowers were then returned to the arena for the start of the next foraging bout.

### Bee trials

Differential conditioning techniques were used to test for the ability of bees to learn the presence of a sucrose reward associated with both humid and dry flower variants in each of the environmental treatment groups as an indicator of floral humidity signal detectability. Individual bees were used just once across all tests, with a record of used bees being kept to avoid repeated use. Bees came from six different nests, which were used on a rotating basis. Bees were assigned to one of the four environmental humidity treatments as described above. Within each of the environments, each bee was also allocated to one of three test groups, which were *i*) Control, *ii*) Humid rewards, and *iii*) Dry rewards. In the Control, all flowers were of the dry variant with dry sponges, meaning there was no humidity difference between rewarding and non-rewarding flowers. In the Humid group, the rewarding flowers were humid variants, and non-rewarding flowers were dry variants. In the Dry group, the rewarding flowers were dry flower variants, and the non-rewarding flowers were humid variants. The feeding wells of flowers held a 25ml of 30% sucrose solution or of water depending on if they were rewarding or non-rewarding, respectively. A summary of the twelve total test groups is shown in Table 1. Twelve bees were used for each test group, and no bee was used more than once, meaning that 144 bees were used in total.

**Table 1.** Summary of the treatment conditions in each environment and test group

| Environmental treatment | Test group |  |  |
| --- | --- | --- | --- |
|  | Control | Humid Rewards | Dry Rewards |
| Low<br>(RH 20-25%) | Rewarding – Dry<br>Non-rewarding – Dry | Rewarding – Humid<br>Non-rewarding – Dry | Rewarding – Dry<br>Non-rewarding – Humid |
| Ambient<br>(RH 35-45%) | Rewarding – Dry<br>Non-rewarding – Dry | Rewarding – Humid<br>Non-rewarding – Dry | Rewarding – Dry<br>Non-rewarding – Humid |
| Medium<br>(RH 67.5-72.5%) | Rewarding – Dry<br>Non-rewarding – Dry | Rewarding – Humid<br>Non-rewarding – Dry | Rewarding – Dry<br>Non-rewarding – Humid |
| High<br>(RH 92.5-97.5%) | Rewarding – Dry<br>Non-rewarding – Dry | Rewarding – Humid<br>Non-rewarding – Dry | Rewarding – Dry<br>Non-rewarding – Humid |

During a learning trial, individual foragers were released alone into the arena, where they would encounter four rewarding and four non-rewarding flowers. The humidity produced by the internal components of the flowers were varied according to the test group that the bee was in, and variants were recognisable to the experimenter by the odd or even numbers on the sides. Once released into the flight arena, the bee was allowed to forage on any flower. Observations were made on two foraging behaviours: *i*) ‘land’, defined as the bee making physical contact with the top of the flower surface at any location even if the bee continued flying, and *ii*) ‘feed’, defined as the bee extending its proboscis into the feeding well. Consequently, each visit could be categorised either as *i*) ‘correct’ where a bee would land and feed from a rewarding flower, or land without feeding from a non-rewarding flower, and *ii*) ‘incorrect’ where a bee would land and feed from a non-rewarding flower or land without feeding from a rewarding flower. These classifications reflect those used in previous conditioning studies (*e*.*g*. Lawson et al. 2018). During each foraging bout, each visit was observed and recorded as either correct or incorrect. At the end of a foraging bout the bee was allowed to return to the nest, and then released back into the arena at the start of the next bout. This was repeated until the bee had completed a total of 70 individual visits to flowers. After each visit, the flower that the bee had landed on was repositioned whether the bee had fed or not to prevent the bee from learning the locations of rewarding and non-rewarding flowers. When a bee departed, the flower was removed by the experimenter through one of the six doors. If the bee had fed from the feeding well, the well was topped up with a fresh 25ml drop of sucrose or water as appropriate. The flower was replaced at a different location in the arena using the doors while the bee was visiting another flower. Any return visits made before a flower had been repositioned and refilled by the experimenter were not counted.

At the end of each foraging bout, all flowers were removed and cleaned as described previously before being returned before the start of the next bout, and all feeding wells cleaned and refilled as described previously. Atmospheric relative humidity conditions were also managed at the end of each bout as described previously. A foraging bout would only begin once all flowers were cleaned and refilled, and the arena humidity was within the defined parameters.

### Statistical analysis

A general linear model (GLM) was constructed in *R* 4.0.5 considering treatment, environment and time period as explanatory variables for the number of correct observations in a ten-visit block, with individual as an error term. Initial exploration (conducted using *SPSS* 26) showed that assumptions of sphericity were violated with the raw number of correct observations, but squaring this value avoided this violation, and inspection of the residuals for each of the separate time periods showed they were sufficiently normal to satisfy model assumptions. Because the analysis considered multiple timepoints, we also checked the assumption that the variance:covariance matrix had compound symmetry, such that all the time gaps between the observations were equally correlated: we did this by rewriting the model in the form of a generalised least squares model, and tested whether assuming that the correlation matrix was symmetrical was sufficient, or whether other correlation assumptions gave a more parsimonious fit to the data. We found that assuming an ARIMA structure to the correlation matrix gave a slightly better fit (measured using AIC) than the symmetrical correlation matrix assumed for the original GLM, but continue to report the initial GLM here as the results were qualitatively identical between the different models and it is simpler to decompose the GLM in order to explore the interactions.

## RESULTS

The number of correct responses made during the ten-visit periods was dependent upon both environmental treatment, rewarding flower type and the time interval (three-way interaction *F*_36, 792_ = 2.41, *p* < 0.005, full model results reported in the supplementary information). Because the three-way interaction was significant, the interactions were explored visually to investigate how best to explore this interaction. It looked most appropriate to decompose the model by the environmental treatment, and test the two-way interactions within each environmental treatment using simple main effects (details in supplementary information). If we do this, each of the two-way interactions between time interval and rewarding flower type is still significant (using Bonferroni corrections to adjust, low humidity environment: *F*_2,792_ = 45.38, *p* < 0.001; ‘ambient’ humidity environment: *F*_2,792_ = 52.45, *p* < 0.001; medium humidity environment: *F*_2,792_ = 50.65, *p* < 0.001; high humidity environment: *F*_2,792_ = 24.21, *p* < 0.001). It makes little biological sense to decompose the two-way interactions further, so we explore the results split by environment here.

In all of the environments, the bees faced with control flowers (where both rewarded and unrewarded flowers were dry, the blue solid lines in figure 1) remained close to the ‘random’ choice line (the thick grey dotted line on the panels of figure 1), suggesting that they were unable to discern the reward before landing and probing the feeding wells. When the bees were challenged with differentiating between humid and dry flowers, they demonstrated that they were able to learn the rewarding flower type in most of the environments, shown by the increase in correct choices over time when either the dry (black dotted line) or humid (orange dashed line) flowers were rewarding, with both flower types being learnt at similar rates, although the dry rewarding flowers appeared to be learnt faster in all the environments. However, in the high humidity environment, the bees appeared unable to learn that humid flowers were rewarding (figure 1d), whilst in the medium humidity environment (with humidity levels above that experienced by the bees outside of the experimental period), the bees appeared to learn that humid flowers could be rewarding at a lower rate than they showed for dry flowers (figure 1c).

**Figure 1.**
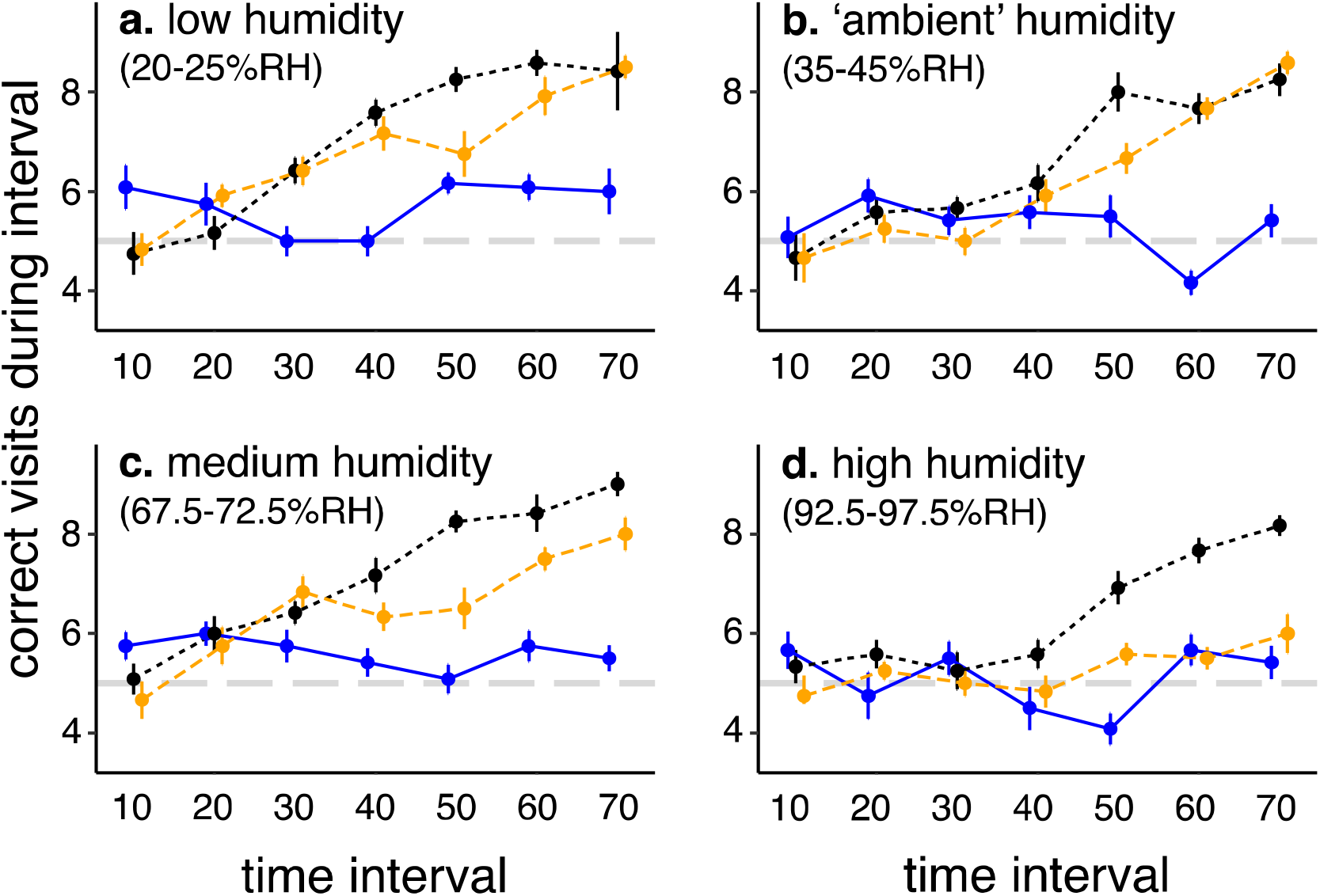
Mean correct visits (±SE) made by *Bombus terrestris* in a ten-visit interval, for bees presented with *a*) low, *b*) ambient, *c*) medium, or *d*) high humidity environments. Blue solid lines represent the control treatment, orange dashed lines represent the rewarding humid flower treatment, and black dotted lines represent the rewarding dry flower treatment.

## DISCUSSION

Signals produced by plants to attract pollinators may be vulnerable to being masked by background noise created by climatic heterogeneity (Candolin 2003, Bradbury and Vehrencamp 2011). If floral humidity signals produced by flowers (von Arx et al. 2012, Harrap et al. 2020) are not robust to environmental humidity fluctuations, then their use in pollination interactions may be limited. This study explored the detectability to bumblebees of a floral humidity signal generated at 2% contrast to ambient conditions when presented against varying levels of background noise in a series of differential conditioning experiments, mimicking the levels of floral humidity that would be experienced by a visiting bumblebee. If floral humidity is robust to environmental fluctuations, it was expected that there would be no effect of increasing noise on the learning response in bees. We found that our artificial humidity signal was detectable and robust to noise as learning was observed across all environments, to varying degrees. Learning was faster when elevated floral humidity was presented as a nonrewarding stimulus, or at greater contrast to the background in in an environment with low humidity. The results also show that the capacity of elevated floral humidity that could be used as a positive stimulus by bees could be constrained by background humidity; lower environmental humidity led to learning occurring, while at high humidity there was no response.

We found that in high humidity environments, the bees had trouble learning that flowers presenting humid signals were rewarding. In the context of high humidity or recent rainfall pollinators may still use floral humidity to accurately determine profitability, but process humidity as a negative stimulus as has been demonstrated by pollinators switching to flowers with concentrated nectar that would be relatively drier (Willmer 1986, Raguso and Willis 2005, Contreras et al. 2013). In this case elevated floral humidity would be a by-product of environmental interference of rewards and no longer be a floral signal. Our observations in high humidity support this view as bees used humidity differences to determine rewards, but were unable to learn a positive association to humid variants. Observations by Harrap et al. (2020, 2021) that floral humidity is produced at varying and sometimes negligible levels, and that bumblebees learnt to associate rewards with both dry and artificial flowers led to the suggestion that floral humidity may in some cases be used instead to confer the identity of a flower to pollinators instead of reward status. The fact that bees learned to associate humidity with both non-rewarding and rewarding flowers supports the interpretation that in some cases floral humidity may be used to confer identity instead of reward status (Harrap et al. 2021), although this is less likely to be the case in extreme conditions, and more likely in conditions of intermediate humidity where osmotic demands are not strongly weighted in either direction. An alternative interpretation may be that bees can be conditioned to learn floral humidity as either a positive or nonrewarding stimulus in ambient conditions simply because they hold the capacity for either response as is required to respond appropriately to humid flowers in varying climatic conditions. In summary, while floral humidity clearly holds capacity to be a useful component in the multimodal display, the relative value of its presence or absence to plants as a signalling strategy depends on the background humidity at a given time.

For a signal to be detectable it must be generated at a level of contrast against the background which meets or exceeds the minimum response threshold of the receiver’s sensory channels, at which point a response is induced (Wiley 2006). Masking occurs when background noise is elevated to an intensity at which the reduction in contrast between a signal and the background means signals are no longer distinguishable, but a robust signal persists amidst noise. Robustness and the noise required for masking are therefore determined by the investment in signal generation by the sender and in sophisticated sensory equipment by the receiver (Wiley 2006). The current study shows that a floral humidity signal produced at 2% contrast to ambient conditions was robust as it was detectable by foraging bees when background noise was elevated to both 72.5-79.5% and 92.5-97.5% RH, where it is expected that signalling contrast was reduced due to the lowering of the osmotic gradient between humid flowers and the surrounding air preventing evaporation (Corbet 1990, Azad et al. 2007, von Arx et al. 2012). The results indicate that despite the reduction in the osmotic gradient, the effect of the environment was not sufficient to completely eliminate the signal from humid artificial flowers, which must have still been producing a floral RH signature elevated above background conditions in the Medium and Humid environments in order to be detected. However, the actual RH value of humid flowers was not measured after initial setup, so the exact extent to which signal contrast was reduced below 2%, if at all, cannot be identified. The robustness of floral humidity will also be partly determined by the sensitivity of the insect hygroreceptor system. In the honeybee *Apis mellifera*, firing activity of dry and moist cells is increased by 1 impulse s^-1^ by instantaneous humidity changes of -2.5% RH and +3.3% RH respectively (Tichy and Kallina 2014). Corresponding observations in the cockroach *Periplaneta americana* were -1.6% RH and + 2% RH, and in the stick insect *Carausius morosus* were -3.3% RH and +2.5% RH (Tichy and Kallina 2014). Sensitivity to the rate of change of humidity is much greater. Changes of 1 impulse^-1^ were induced by altering the rate of humidity change by -0.18% RH s^-1^ and +0.19% RH s^-1^ (Tichy and Kallina 2014). Corresponding observations in the cockroach were and -0.2% RH s^-1^ and +0.1% RH s^-1^, and for the stick insect were -0.4% RH s^-1^ and +0.4 RH s^-1^ (Tichy and Kallina 2014). Furthermore, observations of low sensory resolution in dry or ambient conditions and heightened sensitivity of responses as conditions approach humid extremes have been recorded in the case of mosquitoes *Culex quinquefasciatus*, mealworms *Tenebrio molitor* and human louse *Pediculus humanus* (Thomson 1938, Pielou and Gunn 1940, Wigglesworth 1941), and may help to explain observation in the honeybee (Tichy and Kallina 2014). A response to instantaneous humidity appears unlikely to explain the detectability below 2% unless sensitivities in the bumblebee are also heightened with increasing environmental humidity. However, greater resolution to qualitative change suggests that bees could have responded to even a heavily reduced rate of change of humidity as they entered the floral headspace of humid flowers, encountering a gradient of increasing intensity with proximity to the surface (Harrap et al. 2020).

Where contrast is increased against a background due to a decrease in noise or intensity of production, detectability is enhanced (Lunau et al. 1996) which increases the likelihood of a response in the receiver (Wiley 2006). When background noise was reduced to 20-25% RH, a steeper osmotic gradient would have been established between floral humidity and the background, enhancing evaporative effects and providing a steeper contrast against the resulting signal. The observations of faster learning responses in both groups in this environment support previous studies that emphasise the role of contrasting floral signals in promoting learning and floral constancy. Bumblebees took longer to make decisions and identify rewarding flowers amongst nonrewarding alternatives when presented at low contrast (Spaethe et al. 2001, Chittka and Spaethe 2007, Dyer et al. 2008), and Goulson (2000) reported that foraging bumblebees took twice as long to locate flowers that were presented alongside similarly coloured alternatives than when alone. Furthermore, increasing contrast through colour distance was shown to enhance the speed of developing constancy and the accuracy of decisions (Spaethe et al. 2001, Dyer and Chittka 2004b).

Reduced contrast of olfactory signals had detrimental effects of learning time of bumblebees (Lawson et al. 2017a) and reduced efficiency of reward location in hawkmoths *Manduca sexta* (Cardé and Willis 2008). The observations presented here may also be partly understood by considering the variable sensitivity of hygroreceptor cells, where experiments on the honey demonstrated greater sensitivity when amplitudes of humidity oscillation were increased (Yokohari et al. 1982, Tichy and Kallina 2014). An increased signal contrast may have induced a similar shift in hygroreceptor sensitivity, promoting a faster neural response in low conditions than in other environments. In light of the literature presented, it is not surprising that elevating signal contrast in the current study promoted a faster onset of learning floral humidity differences, whether as a positive or negative association.

The local climate dictates the behavioural responses towards individual flowers and stimuli by regulating demands and preferences of foragers, and by changing floral reward status (Corbet 1990, Chown et al. 2011). The results of the current study show that environmental humidity affected the learning response of bees to rewards associated with humid flowers, accelerating learning in dry conditions but mitigating the response in high humidity. In the natural habitat, relative foraging preferences are likely determined by homeostatic water balances induced partly by environmental humidity (Prange 1996), with aridity promoting water deficit both of colonies and foragers (Willmer 1986, Human et al. 2006, Ellis et al. 2008). For example, bees initiated water collection trips (Willmer 1988) and hawkmoths switched to preferences for more dilute nectar rewards in arid conditions (Contreras et al. 2013). As osmotic demand becomes more weighted, the perceived value of floral humidity signals may increase (von Arx et al. 2012) and floral humidity may become a significant attractant (Willmer 1986). For example, the diluted nectar by the desert dwelling *Echium wildpretii* is thought to function as a water reward to foraging honeybees (Olesen 1988). Conversely, in high humidity energetic demands become more weighted and preferences shift to more concentrated nectar sources (Raguso and Willis 2005). Mason bees switched to concentrated nectar sources in higher humidity (Willmer 1986). In extreme conditions, high water burden may reduce the perceived profitability of humid flowers further still (Bertsch 1984, Willmer 2011). In the natural environment, climatic humidity also affects the reward status of nectar sources. Aridity lowers reward volume by reducing the secretion (Bertsch 1983, Nicolson 1993) and accelerating the evaporation of nectar, which may also reduce palatability by increasing viscosity (Corbet et al. 1979, Plowright 1987, Willmer 2011). Therefore, the perceived profitability of humid flowers is likely to increase in arid conditions. On the other hand, high humidity may cause excessive dilution of nectar which reduces energetic content (Corbet and Delfosse 1984, Cnaani et al. 2006, Willmer 2011). Dilution may be induced by physical contamination of nectar during rainfall, which is, notably, a likely context in which pollinators will experience high humidity extremes (Lawson and Rands 2019). As bee-pollinated flowers tend to hold higher sugar concentrations, diluted nectar is a likely deterrent indicating unfavourable rewards and so the perceived profitability of humid flowers will decrease (Willmer 2011). In both instances, the mechanistic link between nectar and humidity (von Arx 2013, Harrap and Rands 2021) may provide the means by which transient profitability status is assessed. The effects of climate on pollinator preferences and reward status may mean that whilst in arid conditions a floral humidity signal encourages visitation and enhances learning responses towards humid flowers, the same signal in a humid environment may discourage visitation and inhibit floral humidity being learnt as a positive stimulus. Our results appear to mimic this response which is surprising because the bees were only briefly exposed to different humidity conditions when entering the arena to forage. It is unlikely they were exposed long enough for significant shifts in individual osmotic demands to develop before returning to the colony, which itself remained in ambient conditions so would not have contributed to the response. Furthermore, the reward status of artificial flowers was controlled and nectar rewards were not exposed to conditions long enough for equilibration to occur as in natural flowers. Our observations may therefore reflect an innate shift in the perceived profitability of elevated floral humidity induced by the environment which was induced in bees despite no change in actual profitability of humid flowers. In the natural environment, such a shift in innate preference may be useful for quickly avoiding unprofitable flowers and maintaining osmotic balance as conditions change without incurring the time costs associated with learning. It would be interesting to explore how bees and other pollinators responded to natural differences in environmental humidity in the wild, as the natural responses of pollinators may well be dependent upon conditions which could be vastly altered by changes in rainfall and temperature induced by climate change.

## ACKNOWLEDGEMENTS

Heather Whitney, Michael Harrap and Sverre Tunstad are thanked for discussion and assistance in the laboratory.

## AUTHOR CONTRIBUTIONS

ASH and SAR designed the experiment. ASH conducted the experimental work, which was analysed by both authors. The initial draft was written by both authors, based on the thesis produced by ASH. Both authors edited and approved the final version of the manuscript.

## SUPPLEMENTARY DATA

Two files are included:

- The dataset ‘data_in_10s.txt’ is a tab-delimited text file, where ‘environment’ refers to the four environment types (‘low’, ‘ambient’, ‘medium’ or ‘high’), treatment refers to the treatment type (‘control’, ‘dry’ = dry flower reward, ‘humid’ = humid flower rewarding), period refers to the last period of the ten-visit block that the statistic refers to, ‘correct’ is the number of correct visits made during the ten-visit period, and ‘individual’ is the identity of the bee being recorded.
- The code file ‘Harrison_Rands_code.R’ is a text file containing the *R* code used for processing the dataset.

