## Supplementary material for "The ability of bumblebees *Bombus terrestris* (Hymenoptera: Apidae) to detect floral humidity is dependent upon environmental humidity": R code

tens <- read.table("data_in_10s.txt", header=T)
tens$environment <- as.factor(tens$environment)
tens$treatment <- as.factor(tens$treatment)
tens$period <- as.factor(tens$period)
tens$individual <- as.factor(tens$individual)

#I originally ran this as an untransformed model, but this didn't fit assumptions about data sphericity (I checked this using SPSS), so I
#reran the model with the dataset squared. This satisfied sphericity assumptions, and inspection of the standardised residuals produced for
#the seven separate timeperiods showed data were suitably normal.
tenmod1 <- aov(data=tens, correct^2 ~ environment * treatment * period + Error(individual))
summary(tenmod1)

#this gives

# Error: individual
# Df Sum Sq Mean Sq F value   Pr(>F)    
# environment             3  23122    7707  26.773 1.37e-13 ***
#   treatment               2  55396   27698  96.214  < 2e-16 ***
#   environment:treatment   6   5434     906   3.146  0.00648 ** 
#   Residuals             132  38000     288                     
# ---
#   Signif. codes:  0 ‘***’ 0.001 ‘**’ 0.01 ‘*’ 0.05 ‘.’ 0.1 ‘ ’ 1

# Error: Within
# Df Sum Sq Mean Sq F value   Pr(>F)    
# period                         6  90988   15165  86.596  < 2e-16 ***
#   environment:period            18  10446     580   3.314 4.15e-06 ***
#   treatment:period              12  59193    4933  28.168  < 2e-16 ***
#   environment:treatment:period  36  15176     422   2.407 1.05e-05 ***
#   Residuals                    792 138694     175                     
# ---
#   Signif. codes:  0 ‘***’ 0.001 ‘**’ 0.01 ‘*’ 0.05 ‘.’ 0.1 ‘ ’ 1

#note - only the three-way interaction is reported in the manuscript

#I was concerned that the data may not fit assumption of compound symmetry, meaning that the correlations between separate timeperiods were not the same.
#I tested this by rewriting the model as a GLS, and exploring different correlation functions...

#the following uses gls to test whether the assumptions about different time steps having equal variances between them is sensible,
#by using the assumption of compound symmetry. I've followed the example here
#see https://online.stat.psu.edu/stat510/lesson/10/10.1
#by testing different gls models with different assumptions about the symmetry of the correlation matrix for measurements at the seven time points
#the first is the symmetrical case (which is what the earlier model considers), the second makes no assumptions about symmetry, and the third and fourth do similar
#but include an assumption about autocorrelation between consecutive timepoints, with the last being a general ARIMA structure

library(nlme)

tmglsSYMM <- gls(correct^2 ~ environment * treatment * period,data=tens,  corr=corCompSymm(form= ~ 1 | individual))
tmglsNOSTRUCT <- gls( correct^2 ~ environment * treatment * period,data=tens,  corr=corSymm(form= ~ 1 | individual), weights = varIdent(form = ~ 1 | period))
tmglsAR1 <- gls( correct^2 ~ environment * treatment * period, data=tens, corr=corAR1(form= ~ 1 | individual))
tmglsAR1HET <- gls(correct^2 ~ environment * treatment * period, data=tens, corr=corAR1(form= ~ 1 | individual), weights=varIdent(form = ~ 1 | period))
#the following calculates the AIC etc between these, and demonstrates that the symmetrical model is a good fit
anova(tmglsSYMM,tmglsNOSTRUCT, tmglsAR1, tmglsAR1HET)
anova(tmglsSYMM, tmglsAR1HET)
#however this last demonstrates that a general ARIMA structure gives a better fit.

#the following give the outputs for the ARIMA
anova(tmglsAR1HET)
summary(tmglsAR1HET)

#using this ARIMA structured gls, I'm now looking at the interaction plots to see if there's an obvious pattern to start picking apart the three-way interaction,
library(emmeans)
emmip(tmglsAR1HET, period ~ treatment | environment) #looks like period is the important one describing shape
#so, looking at the two different ways of plotting stuff against period...
emmip(tmglsAR1HET, environment ~ period | treatment) #difficult to interpret
emmip(tmglsAR1HET, treatment ~ period | environment) #this one is nice - the humid treatment line is definitely moving around a lot more
#than the other two, so it looks like it would be good to pick this apart by examining the different treatment types separately
#We condition the model using treatment...
mod.t <- emmeans(tmglsAR1HET, ~ period * environment * treatment)
pairs(mod.t, simple="treatment")

#It makes snese to decompose the mode, given the three-way interaction. It is most intuitive to do this with the
#glm, rather than the gls, as both give qualitatively similar results. We start by splitting the data up...
dryenvt <- subset(tens, tens$environment == "low")
medenvt <- subset(tens, tens$environment == "medium")
ambenvt <- subset(tens, tens$environment == "ambient")
hienvt <- subset(tens, tens$environment == "high")

#it's simplest to work out the F values of the decomposed two-way interactions by hand
#Here, I'm dividing the mean sum of squares for the relevant two-way interaction term by the relevant
#error term from the full 'tenmod1' model
#(for 'individual' error, it is 287.8788 with 132 df,
#for 'within' error, it is 175.1187 with 792 df)
drymod1 <- aov(data=dryenvt, correct^2 ~ treatment * period + Error(individual))
summary(drymod1)
#F value for two-way interaction
7948/175.1187
pf(7948/175.1187,2,792, lower.tail = FALSE)
#gives F(2,792) = 45.38, which gives p < 0.001

medmod1 <- aov(data=medenvt, correct^2 ~ treatment * period + Error(individual))
summary(medmod1)
#F value for two-way interaction
8869/175.1187
pf(8869/175.1187,2,792, lower.tail = FALSE)
#gives F(2,792) = 50.65, which gives p < 0.001

ambmod1 <- aov(data=ambenvt, correct^2 ~ treatment * period + Error(individual))
summary(ambmod1)
#F value for two-way interaction
9185/175.1187
pf(9185/175.1187,2,792, lower.tail = FALSE)
#gives F(2,792) = 52.45, which gives p < 0.001

himod1 <- aov(data=hienvt, correct^2 ~ treatment * period + Error(individual))
summary(himod1)
#F value for two-way interaction
4240/175.1187
pf(4240/175.1187,2,792, lower.tail = FALSE)
#gives F(2,792) = 24.21, which gives p < 0.001

#so the two-way interactions when the three-way model is decomposed into separate treatments are still all significant
#I'm not convinced there is any point statistically in decmoposing these further to consider the model with a single term,
#so I'm not running those interactions here!

#THE FOLLOWING CODE IS INCLUDED TO SHOW HOW THE FIGURE WAS GENERATED, AND IS NOT FULLY ANNOTATED
library(ggplot2)
library(gridExtra)
pd <- position_dodge(3)
#converting period values back to numbers for the sake of plotting
dryenvt$period <- as.integer(dryenvt$period)
ambenvt$period <- as.integer(ambenvt$period)
medenvt$period <- as.integer(medenvt$period)
hienvt$period <- as.integer(hienvt$period)

#following function taken from http://www.cookbook-r.com/Graphs/Plotting_means_and_error_bars_(ggplot2)/
summarySE <- function(data=NULL, measurevar, groupvars=NULL, na.rm=FALSE,
                      conf.interval=.95, .drop=TRUE) {
  library(plyr)
  
  # New version of length which can handle NA's: if na.rm==T, don't count them
  length2 <- function (x, na.rm=FALSE) {
    if (na.rm) sum(!is.na(x))
    else       length(x)
  }
  
  # This does the summary. For each group's data frame, return a vector with
  # N, mean, and sd
  datac <- ddply(data, groupvars, .drop=.drop,
                 .fun = function(xx, col) {
                   c(N    = length2(xx[[col]], na.rm=na.rm),
                     mean = mean   (xx[[col]], na.rm=na.rm),
                     sd   = sd     (xx[[col]], na.rm=na.rm)
                   )
                 },
                 measurevar
  )
  
  # Rename the "mean" column    
  datac <- rename(datac, c("mean" = measurevar))
  
  datac$se <- datac$sd / sqrt(datac$N)  # Calculate standard error of the mean
  
  # Confidence interval multiplier for standard error
  # Calculate t-statistic for confidence interval: 
  # e.g., if conf.interval is .95, use .975 (above/below), and use df=N-1
  ciMult <- qt(conf.interval/2 + .5, datac$N-1)
  datac$ci <- datac$se * ciMult
  
  return(datac)
}

drysumm <- summarySE(dryenvt, measurevar="correct", groupvars=c("period","treatment"))
ambsumm <- summarySE(ambenvt, measurevar="correct", groupvars=c("period","treatment"))
medsumm <- summarySE(medenvt, measurevar="correct", groupvars=c("period","treatment"))
hisumm <- summarySE(hienvt, measurevar="correct", groupvars=c("period","treatment"))

ggdry <- ggplot(drysumm, aes(x=period, y=correct, colour=treatment))+
  geom_hline(yintercept=5, linetype="longdash",colour="grey85", size = 1) + 
  theme_classic()+
  geom_errorbar(aes(ymin=correct-se, ymax=correct+se), width=.1, position=pd) +
  geom_line(position=pd, aes(linetype=treatment))+
  geom_point(position=pd)+
  scale_color_manual(values=c("blue","black","orange"))+
  theme(legend.position = "none")+
  ylim(3.5,9.5)

  
ggamb <- ggplot(ambsumm, aes(x=period, y=correct, colour=treatment)) + 
  geom_hline(yintercept=5, linetype="longdash",colour="grey85", size = 1) + 
  theme_classic()+
  geom_errorbar(aes(ymin=correct-se, ymax=correct+se), width=.1, position=pd) +
  geom_line(position=pd, aes(linetype=treatment)) +
  geom_point(position=pd)+
  scale_color_manual(values=c("blue","black","orange"))+
  theme(legend.position = "none")+
  ylim(3.5,9.5)
ggmed <- ggplot(medsumm, aes(x=period, y=correct, colour=treatment)) + 
  geom_hline(yintercept=5, linetype="longdash",colour="grey85", size = 1) + 
  theme_classic()+
  geom_errorbar(aes(ymin=correct-se, ymax=correct+se), width=.1, position=pd) +
  geom_line(position=pd, aes(linetype=treatment)) +
  geom_point(position=pd)+
  scale_color_manual(values=c("blue","black","orange"))+
  theme(legend.position = "none")+
  ylim(3.5,9.5)
gghi <- ggplot(hisumm, aes(x=period, y=correct, colour=treatment)) + 
  geom_hline(yintercept=5, linetype="longdash",colour="grey85", size = 1) + 
  theme_classic()+
  geom_errorbar(aes(ymin=correct-se, ymax=correct+se), width=.1, position=pd) +
  geom_line(position=pd, aes(linetype=treatment)) +
  geom_point(position=pd)+
  scale_color_manual(values=c("blue","black","orange"))+
  theme(legend.position = "none")+
  ylim(3.5,9.5)

grid.arrange(ggdry, ggamb, ggmed, gghi, ncol=2)


ggplot(drysumm, aes(x=period, y=correct, colour=treatment))+
  geom_hline(yintercept=5, linetype="longdash",colour="grey85", size = 1) + 
  theme_classic()+
  geom_errorbar(aes(ymin=correct-se, ymax=correct+se), width=.1, position=pd) +
  geom_line(position=pd, aes(linetype=treatment))+
  geom_point(position=pd)+
  scale_color_manual(values=c("blue","black","orange"))+
  ylim(3.5,9.5)
